# PlasEval: a framework for comparing and evaluating plasmid detection tools

**DOI:** 10.1101/2024.04.30.591963

**Authors:** Aniket Mane, Haley Sanderson, Aaron P. White, Rahat Zaheer, Robert G. Beiko, Cedric Chauve

## Abstract

Plasmids play a major role in the transfer of antimicrobial resistance (AMR) genes among bacteria via horizontal gene transfer. The identification of plasmids in short-read assemblies is a challenging problem and a very active research area. *Plasmid binning* aims at detecting, in a draft genome assembly, groups (bins) of contigs likely to originate from the same plasmid. Several methods for plasmid binning have been developed recently, such as PlasBin-flow, HyAsP, gplas, MOB-suite, and plasmidSPAdes. This motivates the problem of evaluating the performances of plasmid binning methods, either against a given ground truth or between them.

We describe PlasEval, a novel method aimed at comparing the results of plasmid binning tools. PlasEval computes a dissimilarity measure between two sets of plasmid bins, that can originate either from two plasmid binning tools, or from a plasmid binning tool and a ground truth set of plasmid bins. The PlasEval dissimilarity accounts for the contig content of plasmid bins, the length of contigs and is repeat-aware. Moreover, the dissimilarity score computed by PlasEval is broken down into several parts, that allows to understand qualitative differences between the compared sets of plasmid bins. We illustrate the use of PlasEval by benchmarking four recently developed plasmid binning tools – PlasBin-flow, HyAsP, gplas, and MOB-recon – on a data set of 54 *E. coli* bacterial genomes.

PlasEval is freely available at https://github.com/acme92/PlasEval

## 1 Introduction

Mobile Genetic Elements (MGEs) are DNA sequences within genomes that have the ability to move or be transferred within and between bacterial cells; MGEs can carry important genes, such as virulence factors or antimicrobial resistance (AMR) genes [23, 13]. Plasmids form an important family of MGEs due to their ability to transfer between bacteria of different species through horizontal gene transfer, thus contributing to the spread of AMR genes. Due to their high mobility and the function of the genes they carry, detecting plasmids in bacterial isolates is important, whether it is for ecological studies or motivated by public health (e.g. tracking the spread of AMR genes in an infectious disease outbreak) [27]. This has motivated the development of bioinformatics tools to identify plasmids from draft bacterial genomes, a process called *plasmid binning*. Plasmid binning aims to detect, from a draft genome assembly, groups of contigs (called *plasmid bins*) that are assumed to originate from the same plasmid that was present in the sequenced isolate(s). Plasmid binning is known to be a challenging problem [6] and is an active research area [3, 25, 22, 4, 20, 19].

With multiple plasmid binning methods developed to-date, choosing which one to apply in a given project for which plasmids are of interest is far from obvious. Suitability of a method can only be validated by applying various considered methods on a test data set, for which the true plasmids are known. Each true plasmid is represented by its corresponding plasmid bin harbouring the set of short read contigs that are mapped to it. These plasmid bins are hereafter referred to as the *ground truth*.

Such a data set can be built in several ways. It can be generated by analyzing isolates for which both closed genome assemblies, including plasmids, and sequencing data are publicly available, for example in public databases such as RefSeq and GenBank for closed assemblies and SRA for sequencing data. These testing samples can then be re-assembled into draft assemblies, after which ground truth plasmid bins can be obtained from mapping the resulting contigs against the true plasmids. Alternatively, isolates for which hybrid sequencing data, composed of both short reads (e.g. Illumina data) and long reads (e.g. Oxford Nanopore data), can be used. Hybrid assemblies tend to be more amenable to detecting plasmids than short-read or long-read assemblies individually [11, 18, 8]; once hybrid contigs identified as plasmids have been obtained, they can be used as in the previous case to define ground truth plasmid bins (see [30] for example). Nevertheless, given the plasmid bins predicted by a plasmid binning tool and ground truth plasmid bins, it is essential to assess the accuracy of the predicted plasmid bins through well-defined accuracy measures. Additionally, it is also of interest to compare the predictions of different plasmid-binning tools for the same assembly in order to identify plasmids that are consistently predicted across all the tools.

Accuracy measures developed to assess the quality of full genome assemblies [21] are not fully relevant for the plasmid binning problem. This has motivated the development of accuracy measures specific to the plasmid binning problem. The recent papers introducing gplas [4] and PlasBin-flow [19] introduced statistics inspired by the classical precision, recall and F1-score used to evaluate classification and clustering methods, that provide a high-level measure of accuracy. However, such high-level statistics do not immediately provide insight into the specific errors made by different methods; for example, before deciding which plasmid binning method to use, it might be of interest to know if incorrectly predicted plasmid bins result from true plasmid bins being split into several predicted bins or from the mixing of several true plasmid bins into several predicted plasmid bins. Moreover, comparing plasmid binning methods in absence of a ground truth, for example to understand the nature of the differences in the plasmid bins predicted by the considered tools could be valuable.

Here we introduce a novel measure of dissimilarity between a pair of sets of plasmid bins for a given draft genome assembly, implemented in the PlasEval software. The dissimilarity measure accounts for the difference in terms of contig content between two sets of plasmid bins, weighted by contig lengths. Mathematically, it is defined in terms of a parsimonious scenario of bin *splits* (contigs from a plasmid bin being split into two bins), followed by *joins* (two plasmid bins joined to form a single bin). The definition of the dissimilarity using splits and joins allows to separate the dissimilarity score between two sets of plasmid bins (either a set of predicted bins and a corresponding set of ground truth bins, or two sets of predicted bins) into different parts that provide both qualitative and quantitative insight into the nature of the differences between the two sets of bins. In Section 3, we introduce this dissimilarity measure together with an algorithm to compute it. We also describe practical cases in which the PlasEval tool is of interest. Section 4 describes the results of our experiments, using PlasEval to evaluate four plasmid binning methods (MOB-recon [25], HyAsP [22], gplas [4] and PlasBin-flow [19]) on a data set of 54 *Escherichia coli* samples for which hybrid sequencing data is used to define ground truth plasmid bins. Our experiments illustrate the added value of PlasEval compared to high-level statistics such as precision and recall.

## 2 Preliminaries

Plasmids are extra-chromosomal molecules present in a bacterial cell; they are generally much shorter than the chromosome(s), ranging from a few hundred nucleotides to less than a megabase, with larger plasmids being rare [29]. Most plasmids are circular molecules, and they can occur within a cell in multiple copies, up to hundreds of copies, although larger plasmids tend to be single-copy. Plasmids often harbor repeats (such as insertion sequences), that can be repeated within a plasmid, between plasmids or between plasmids and the chromosome.

*Plasmid binning*. Given a contig assembly with contig set *𝒞*, a *plasmid bin P is a set of contigs* (i.e. is an unordered subset of *𝒞*). The *plasmid binning* problem aims to compute, from the contig assembly of a bacterial isolate, a set of plasmid bins, ideally representative of the plasmid content of the isolate (i.e. where the contigs in each plasmid bin are the contigs of a plasmid of the isolate).

Due to the frequent presence of repeats in plasmid, a given contig *c* ∈ *𝒞* can appear in several plasmid bins. A contig can also be repeated within a plasmid (which would make a plasmid bin a multiset of contigs), but most plasmid binning methods do not account for such features and we do not consider this case here. Plasmid binning from a short-reads contigs assembly is a challenging problem [6]; advances in long-reads sequencing technologies allow to detect plasmids with greater accuracy [29] although they often miss shorter plasmids [17]. Nevertheless, in a practical context such as epidemiological surveillance, standard approaches still rely on short-reads sequencing [14], thus we focus this work on evaluating plasmid binning methods from a short-reads contigs assembly.

*Plasmid binning methods*. Initial approaches to detect plasmids from an assembly focused on the problem of *contigs classification*. In the contig assembly of a bacterial isolate, a contig is *plasmidic* if the corresponding sequence occurs only in some plasmid(s) of the sequenced isolate, *chromosomal* if it belongs only to the chromosome of the isolate, and *ambiguous* if it is a sequence that occurs both in some plasmid and in the chromosome. Contig classification aims to label contigs as either plasmidic, chromosomal or ambiguous, often using machine-learning approaches (see [30, 5]; [1, 7, 24]). The first plasmid binning methods included PlasmidSPAdes [3] and Recycler [26], both leveraging the information provided by the *assembly graph* (a graph whose vertices are contigs and edges indicate putative contiguity between contigs) to detect plasmid bins as cyclic groups of contigs that satisfy read-depth coverage consistency indicative of multi-copy plasmids. The idea of using the assembly graph, together with plasmid-specific features (presence of known plasmid genes, GC content, read-depth used as a proxy of copy number), is at the core of the most recent plasmid binning methods. Both HyAsP [22] and gplas [4] rely on heuristics to compute plasmid bins as walks in the assembly graphs, while PlasBin [20] and its recent extension PlasBin-flow [19] are based on Mixed Integer Linear Programming to compute plasmid bins as connected subgraphs of the assembly graph. MOB-recon [25] is the only method that does not make use of the assembly graph, relying instead on the comparison of the contigs against a carefully curated database of known plasmids and plasmid-specific genes.

*Measuring plasmid binning accuracy*. Most papers on plasmid binning methods do measure the accuracy of a set of plasmid bins by comparing them to a *ground truth* set of plasmid bins using variations of the notions of precision, recall and F1-score. In our work, we compare the novel measure of accuracy that we introduce to the F1-score defined in [19], that we define formally below.

For a given contig set *𝒞*, let *𝒜* be the set of plasmid bins predicted by a plasmid binning tool and *ℬ* the set of true plasmid bins (ground truth). For a set of contigs *X*, we define the cumulative length of the contigs in *X* as *L*(*X*). For a predicted plasmid bin *P* ∈ *𝒜* and a ground truth plasmid bin *T* ∈ *ℬ*, we define the overlap between *P* and *T* as the cumulative length of the contigs in the intersection of the two sets of contigs *P, T* : overlap(*P, T*) = *L*(*P ∩ T*). The precision and recall of the set *𝒜* of predicted plasmid bins compared to the ground truth *ℬ* are respectively defined as

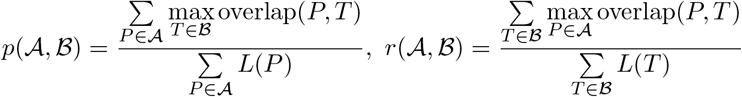

and the F1-score is the arithmetic mean of the precision and recall, 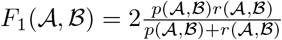

## 3 Methods

The framework we propose aims at computing a weighted set-theoretic dissimilarity measure between two inferred sets of plasmid bins. It is based on the idea of transforming one set of plasmid bins into the other one by a sequence of operations splitting plasmid bins, followed by a sequence of operations joining the resulting splitted bins, while accounting for the contig lengths, unequal contig content and the possible presence of repeated contigs.

In the remainder of this section, we denote by *𝒞* the contig alphabet and consider two sets of plasmid bins denoted by *𝒜* and *ℬ* over *𝒞*. We denote by *D*_*α*_(*𝒜, ℬ*) the dissimilarity between *𝒜* and *ℬ*, where *α* ∈ [0, 1] is a parameter used to weight how contig lengths contribute to the dissimilarity score. We first describe how to compute efficiently *D*_*α*_(*𝒜, ℬ*) in the simpler context where no contig is repeated, then describe how we handle repeated contigs through a branch-and-bound algorithm. We conclude this section by a discussion on theoretical properties of the dissimilarity measure we introduce.

### 3.1 Dissimilarity without repeated contigs

We first consider the case where each contig *c* ∈ *𝒜* appears in at most one plasmid bin in *𝒜* and at most one plasmid bin in *ℬ*. We denote by unique(*𝒜, ℬ*) the set of *unique* contigs *c* such that *c* appears in *𝒜* but not *ℬ* or conversely, and by *𝒜*′ and *ℬ*′ the sets of plasmid bins obtained by deleting respectively from *𝒜* and *ℬ* these unique contigs. For a contig *c*, we denote by *𝓁*(*c*) its length. For a set of contigs *X*, we define *L*_*α*_(*X*) as *L*_*α*_ (*X*) = (∑_*c* ∈ *X*_ ℓ(c))^*α*^, where *α* ∈ [0, 1] is a fixed parameter to weight the contribution of contig lengths to the dissimilarity score.

We first define the operation of splitting and joining plasmid bins, together with the corresponding cost, defined in terms of the lengths of the involved contigs. Figure 1 illustrates these two operations.

**Fig. 1:**
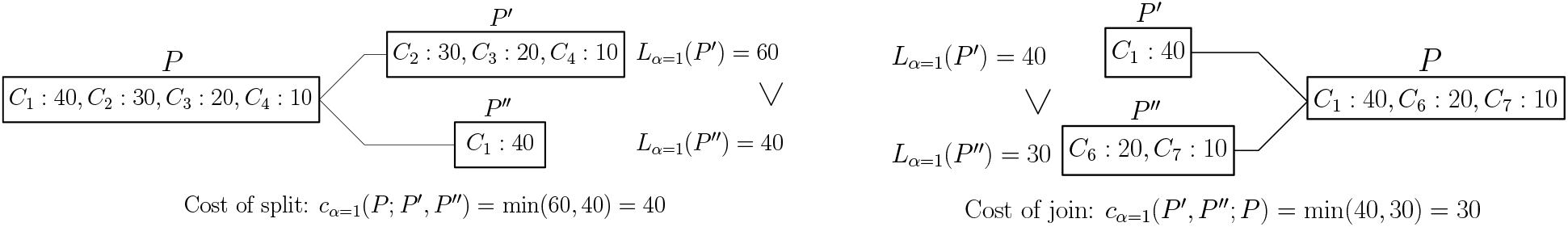
Cost of split and join operations with *α* = 1

#### Definition 1

**(Plasmid bin split and join)**.

*Let P be a set of contigs (plasmid bin); a split of P is composed of two non-empty sets P*′, *P* ′′ *such that P* ′ *∩ P* ′′ = *∅ and P* ′ *∪ P* ′′ = *P*. *The cost of splitting P into P* ′, *P* ′′ *is c*_*α*_(*P* ; *P* ′, *P* ′′) = min(*L*_*α*_(*P* ′), *L*_*α*_(*P* ′′)).

*A join is the symmetric operation: it takes two non-empty sets of contigs P* ′, *P* ′′ *and joins them into a single set P*. *The cost of joining P* ′, *P* ′′ *into P is c*_*α*_(*P* ′, *P* ′′; *P*) = min(*L*_*α*_(*P* ′), *L*_*α*_(*P* ′′)).

The similarity measure *D*_*α*_(*𝒜, ℬ*) we introduce can be defined informally as being composed of two main parts (1) discarding from *𝒜* and *ℬ* their unique contigs, and accounting for the total length of these unique contigs, and (2) transforming *𝒜*′ into *ℬ*^*′*^ by first splitting the plasmid bins of *𝒜*^*′*^ and then joining the resulting bins to obtain *ℬ*^*′*^. Definition 2 below defines the set of bins obtained after the splitting phase.

#### Definition 2.

*For a plasmid bin P in 𝒜*^*′*^, *we define its refinement with respect to ℬ*^*′*^, *denoted by* ref(*P, ℬ*^*′*^), *as the unique partition of P into non-empty sets P*_1_, …, *P*_*k*_ *of contigs such that for any P*_*i*_ *there exists Q* ∈ *ℬ*′ *with P*_*i*_ = *P ∩ Q*.

We denote by ref(*𝒜*^*′*^, *ℬ*^*′*^) the set of plasmid bins obtained by refining all plasmid bins in *𝒜*^*′*^ with respect to *ℬ*^*′*^. It is straightforward to see that ref(*ℬ*^*′*^, *𝒜*^*′*^) = ref(*𝒜*^*′*^, *ℬ*^*′*^). We also denote, for a plasmid bin *P* of *𝒜*^*′*^ (resp. *Q* from *ℬ*^*′*^) by *c*_*α*_(*P* ; *P*_1_, …, *P*_*k*_) (resp. *c*_*α*_(*Q*_1_, …, *Q*_*k*_; *Q*)) the cost of a most parsimonious sequence of splits (resp. joins) creating the bins *P*_1_, …, *P*_*k*_ of ref(*P, ℬ*^*′*^) (resp. *Q*_1_, …, *Q*_*k*_ of ref(*Q, 𝒜*^*′*^)).

#### Definition 3

**(Dissimilarity with no repeated contig)**.

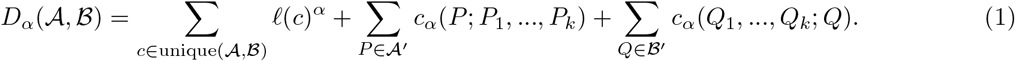

The first term in (1) accounts for unique contigs, while the second term records the cost of splitting *𝒜*^*′*^into ref(*𝒜*^*′*^, *ℬ*^*′*^), and the third term the cost of joining ref(*𝒜*^*′*^, *ℬ*^*′*^) into *ℬ*^*′*^.

We now turn to the cost of refining a set of bins *P* into {*P*_1_, …, *P*_*k*_} as defined in Definition 2. Lemma 1 below, whose proof is straightforward, states that a most parsimonious way to refine a plasmid bin is to perform splits that create the sets *P*_1_, …, *P*_*k*_ by increasing order of size: assuming *L*(*P*_1_) *≤ L*(*P*_2_) *≤ · · · ≤ L*(*P*_*k*_), creating *P*_1_ out of *P* by a single split, then *P*_2_ out of *P − P*_1_ by a split, and so on. By symmetry of splits and joins the same approach also applies to compute *c*_*α*_(*Q*_1_, …, *Q*_*k*_; *Q*).

#### Lemma 1.

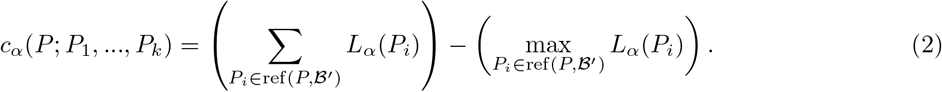

Thus, in the absence of repeated contigs, *D*_*α*_(*𝒜, ℬ*) can be computed easily by the following steps:

– first discarding unique contigs to create *𝒜*^*′*^, *ℬ*^*′*^, which contributes ∑ _*c* ∈ unique(*A,B*)_ *𝓁*(*c*)^*α*^ to *D*_*α*_(*𝒜, ℬ*),
– then splitting *𝒜*^*′*^ into ref(*𝒜*^*′*^, *ℬ*^*′*^), with cost as defined in Lemma 1,
– finally joining ref(*𝒜*^*′*^, *ℬ*^*′*^) into *ℬ*^*′*^, or equivalently splitting *ℬ*^*′*^ into ref(*𝒜*^*′*^, *ℬ*^*′*^), with cost as defined in Lemma 1.

All these steps can be done in polynomial time, so in the absence of repeated contig, *D*_*α*_(*𝒜, ℬ*) can be computed in polynomial time.

### 3.2 Dissimilarity with repeated contigs

To handle the presence of repeated contigs, we follow the *maximal matching* approach used to compute genome rearrangement distances with repeated genes.

Let *c* be a contig that appears in *k*_1_ copies in *𝒜* and *k*_2_ copies in *ℬ*; we call the set of all copies of *c* a *contig family* and we say this family is a *repeat family* if *k*_1_ + *k*_2_ *>* 2. We denote by repeats(*𝒜, ℬ*) the set of all repeat families in *𝒜, ℬ*.

A maximal matching *M*_*c*_ for a contig family *c*, with respectively *k*_1_, *k*_2_ copies of *c* in *𝒜, ℬ*, is composed of *k* = min(*k*_1_, *k*_2_) pairs of contigs *c* (called edges), each pair containing one copy of *c* in each of *𝒜, ℬ*. Given *M*_*c*_, if we label each edge by a unique integer *i* ∈ {1, …, *k*} and rename with *c*_*i*_ both extremities of the edge (contigs initially labelled *c*), while the remaining (if any) unmatched copies of *c* that do not belong to any edge of *M*_*c*_ are labeled arbitrarily *c*_*j*_, *j* ∈ {*k* + 1, …, *k*_1_ + *k*_2_}, we say that the contig family *c* has been *resolved*, i.e. that all copies of *c* have been renamed in such a way that they can be considered as distinguishable.

Let *M* be a set of maximal matchings *M*_*c*_, one for each contig family *c* ∈ repeats(*𝒜, ℬ*); we denote by *M* (*𝒜*), *M* (*ℬ*) the corresponding resolved sets of plasmid bins, where all repeated contigs have been renamed as described above, resulting in an instance with no repeated contig, for which *D*_*α*_(*M* (*𝒜*), *M* (*ℬ*)) can then be defined as in Definition 3 and computed in polynomial time.

#### Definition 4

**(Dissimilarity with repeated contig)**.

*Let ℳ*(*𝒜, ℬ*) *be the set of all sets of maximal matchings resolving all repeat families in 𝒜, ℬ*.

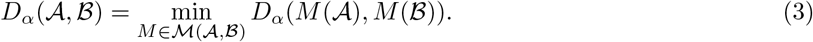

To search the space *ℳ* (*𝒜, ℬ*) of all matchings resolving repeat families in order to find one that minimizes the resulting dissimilarity, we implemented a branch-and-bound algorithm that we describe now at a high level, more details being provided in Appendix B.

The branch-and-bound algorithm is based on exploring a search tree that builds incrementally the sets of bins *𝒜, ℬ* by adding contigs from repeat families, updating the distance as it goes on in a monotonic way.

– The root of the search tree (level 0) is defined as the instance *𝒜*_0_, *ℬ*_0_ obtained by keeping only non-repeat contig families; due to the absence of repeated contigs, *D*_*α*_(*𝒜*_0_, *ℬ*_0_) can be computed in polynomial time.
– Each level *i* of the search tree corresponds to adding the contigs of exactly one repeat family to *𝒜*_*i−*1_, *ℬ*_*i−*1_ in all possible ways, to define larger instances.
– Assume the considered repeat family is for contig *c*, having *k*_1_, *k*_2_ copies in *𝒜, ℬ* respectively. There are 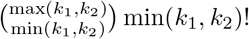 possible maximal matchings between the copies of *c* in *𝒜*_*i*_, *ℬ*_*i*_, and each matching defines a way to add all copies of *c* to *𝒜*_*i−*1_, *ℬ*_*i−*1_ thus creating a larger instance *𝒜*_*i*_, *ℬ*_*i*_ with no repeated contig and for which the dissimilarity *D*_*α*_(*𝒜*_*i*_, *ℬ*_*i*_) can be computed in polynomial time; the set of all these instances form the level *i* of the search tree.
– Finally, a subtree of the search tree, say rooted at level *i*, is explored only if the dissimilarity value *D*_*α*_(*𝒜*_*i*_, *ℬ*_*i*_) of the (partial) instance *𝒜*_*i*_, *ℬ*_*i*_ at its root is strictly lower than the best found dissimilarity value for a full instance; this approach is a proper bounding method as Lemma 2 (Appendix A) shows that adding contigs can never decrease the dissimilarity value.

In some cases, especially for sets of plasmid bins obtained from an assembly with a large number of short contigs, the branch-and-bound algorithm can be time-consuming. To handle such cases, we allow users to set a *minimum length* parameter *𝓁* (default value *𝓁* = 0) and all contigs of length below *𝓁* are discarded from the considered plasmid bins.

### 3.3 Normalized dissimilarity

In order to compare the dissimilarity we introduced to the notions of precision, recall and F1-score, we normalize it between 0 and 1, by dividing it by the sum of the weighted lengths of the contigs appearing in *𝒜* and*ℬ*:

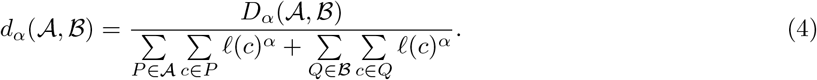

Note that this normalization accounts for the presence of repeats as a contig *c* appearing in several bins will contribute by *𝓁*(*c*)^*α*^ according to it copy number in both *𝒜* and *ℬ*. It is straightforward to prove that (1) *d*_*α*_(*𝒜, ℬ*) = 0 if and only if *𝒜* = *ℬ*, and (2) *d*_*α*_(*𝒜, ℬ*) = 1 if and only if *𝒜* and *ℬ* have no common contig, i.e. every contig appears only in either *𝒜* or *ℬ*.

### 3.4 Discussion

If one considers *𝒜* as a set of plasmid bins predicted by a plasmid binning tool and *ℬ* as the ground truth for the same sample, *D*_*α*_(*𝒜, ℬ*) accounts for four kinds of errors in the predicted bins: contigs that are erroneously predicted as plasmidic (*extra contigs*), plasmidic contigs that are not included in any predicted bin (*missed contigs*), contigs from different true bins that are placed into the same predicted bin (accounted for by splits) and contigs of a true bin that are separated into several predicted bins (accounted for by joins).

The parameter *α* allows to control the contribution of contig lengths in the dissimilarity score. With *α* = 0, the dissimilarity is purely set-theoretic and counts only splits and joins, augmented by 1 for the presence of extra contigs and 1 for missing contigs, and thus does not account at all for contig length. With *α* = 1, the contribution of splits (resp. joins) to the dissimilarity score is exactly the *precision* (resp. *recall*) as defined in [19] (see Section 2). Using *α* = 1 leads to the issue that the fragmentation of a true plasmid bin into several predicted bins is not accounted for. To illustrate this, consider a true plasmid bin *P* with 2*k* contigs, all of the same length, and two sets of predicted bins, *𝒫* _1_ composed of two bins each containing *k* contigs, and *𝒫*_2_ composed of one bin of *k* contigs and *k* bins of one contig each. We then have *d*_1_(*𝒫*_1_, {*P* }) = *d*_1_(*𝒫*_2_, {*P* }) = 0.5, which is the value of both the precision and the recall, thus not accounting for the higher fragmentation of *𝒫*_2_, while for any *α <* 1, *d*_1_(*𝒫*_1_, {*P* }) *< d*_1_(*𝒫*_2_, {*P* }). In our experiments, we use *α* = 0.5.

The core of the definition of *D*_*α*_(*𝒜, ℬ*) is to transform the set of plasmid bins *𝒜*′ into *ℬ*′, where both have equal contig content. This relates to the *syntenic distance* introduced in [15] as a genome rearrangement distance for genomes represented as unordered sets of genes, whose computation is NP-complete [12]. It differs in two major points: (1) while splits and joins correspond to fissions and fusions, our approach does not consider the operation *translocations* considered in the syntenic distance, that would correspond to exchanging sets of contigs between two plasmid bins, and (2) we consider only sequences of splits followed by joins and do not consider mixing both operations. Including translocation in measuring the dissimilarity between sets of plasmid bins would potentially capture more complex errors, such as the mixing of two true plasmid bins into two predicted bins, at the expense of tractability. Considering arbitrary sequences of splits and joins would result in a lower dissimilarity score, although we motivate our choice of considering only sequences of splits followed by joins by the fact that the structure of the intermediate set of bins obtained after all splits, ref(𝒜′, ℬ′), is indicative of the structural differences between the two considered sets of plasmid bins. Moreover, this approach leads to an efficient way to compute the dissimilarity with no repeated contigs, which is crucial in ensuring that the branch-and-bound algorithm that considers repeats does finish in a reasonable computational time.

### 3.5 Practical use of PlasEval

PlasEval is designed to provide computational biologists and bioinformaticians with a precise way to assess the performances of methods aimed at detecting plasmids from short read draft assemblies.

The main application of PlasEval is to compare a set of plasmid binning tools. Given a test dataset of samples for which both short read assemblies and ground truth plasmid bins are available, PlasEval allows to determine the strengths and weaknesses of all considered tools, with high-level statistics (the dissimilarity score) and more refined statistics in terms of the four components of the dissimilarity score (missing and extra contigs, plasmid bins splits and joins). This can be motivated by several applications:

– Comparing a novel plasmid binning tool against a set of state-of-the-art existing plasmid binning tools.
– Investigating a set of plasmid binning tools with the aim to chose one (or several) to apply on a specific dataset.
– Benchmarking a set of plasmid binning tools either against a dataset for which the ground truth plasmid bins are known.

In the first two cases, using PlasEval requires the availability of samples for which the true plasmid bins are known. This can be obtained from samples for which short read assemblies are available or can be computed from sequencing data and either (1) annotated closed genomes are available or (2) hybrid (short and long read) assemblies are available. In Section 4 we describe a simple protocol to determine ground truth plasmid bins from short read assemblies and hybrid assemblies.

## 4 Results

In this section, we use PlasEval to evaluate four plasmid binning tools on a set of 54 samples from a collection of *Escherichia coli* genomes sequenced using both short and long reads by Sanderson et al [28].

### 4.1 Data

The isolates in this analysis belong to a collection of *E. coli* strains collected from Saskatchewan broiler farms by Sanderson et al [28]. The hybrid assemblies, combining short and long reads, were obtained using Unicycler (v0.5.0) [31] in hybrid mode, and were used to define the ground truth for each sample. The hybrid assemblies are available under NCBI BioProject accession number PRJNA912639. The short-read assemblies were generated as follows: Illumina short-read quality was checked using FastQC (v0.11.9) [2], then short reads were trimmed and the adaptors were removed using Fastp (v0.23.4) [10], then the reads were assembled using Unicycler (v0.5.0) [31]. The quality of the short-read assemblies was assessed with Quast (v5.0.2) [16]. For each hybrid assembly, generated hybrid contigs were labelled as chromosome, plasmid or ambiguous based on contig length and circularity. Contigs that were longer than 500, 000 bp were labelled as chromosomes, irrespective of circularity. Circular contigs shorter than 500, 000 bp were labelled as plasmids, while the remaining contigs were labelled as ambiguous. In order to have high-quality ground truth, we selected 54 hybrid assemblies with no ambiguous contigs collectively containing a total of 192 plasmidic contigs, called plasmids from now on. The plasmids had an average length of 48, 657 bp, with the largest plasmid in the dataset being of length 206, 436 bp.

The plasmid binning tools we evaluate are designed to predict plasmid bins from short-read assemblies. We assembled the short reads for the 54 samples using Unicycler, that generates, together with a set of contigs, an *assembly graph* that is used by three of the considered plasmid binning tools. The short-read assemblies of the 54 samples consists of 18, 354 contigs in total: 887 of these contigs are longer than 10^5^ bp, 1, 866 contigs are between 10^4^ bp and 10^5^ bp, 1, 831 are between 10^3^ bp and 10^4^ bp, 7, 901 are between 100 bp and 1, 000 bp, while the remaining 5, 869 contigs are shorter than 100 bp. The average length of the short-read contigs is 15, 033 bp, with the longest contig being of length 790, 743 bp.

The hybrid assemblies and short-read assemblies are available at https://zenodo.org/records/10785151, and at NCBI in the BioProject PRJNA912639. We refer to these genomes here by their NCBI BioSample accessions; for example, “SAMN32247327” is the accession for the genome with the assigned isolate name “EC B1 9226 C5 H CuN CeP” that is accessible at https://www.ncbi.nlm.nih.gov/biosample/SAMN32247327/ in the BioProject.

### 4.2 Experimental setup

For all samples, the plasmid binning tools MOB-recon [25], HyAsP [22], gplas [4] and PlasBin-flow [19] were used on the short-read assemblies to predict plasmid bins. MOB-recon, being a homology-based method, selects contigs potentially belonging to plasmids by mapping them against a plasmid gene database of plasmid replicons and relaxase genes. It also assigns the contig a cluster label. Contigs with the same cluster label are then grouped together to form a plasmid bin. Gplas uses a plasmid identification tool to assign probabilities for each contig belonging to a plasmid or chromosome. It then forms a plasmidome network by identifying plasmid-like walks in the assembly graph. This network is then partitioned into components that represent plasmid bins. HyAsP identifies seed contigs in the assembly graph; these contigs have a high proportion of genes from known plasmid gene databases. It tries to identify circuits or trails in the assembly graph by a greedy approach, aiming to optimize an objective function that accounts for a prior plasmid score for each contig, GC content and depth of sequencing used as a proxy for copy number; the computed circuit or trail define each a plasmid bin. PlasBin-flow follows the general principle of HyAsP but defines plasmid bins as connected subgraphs of the assembly graphs identified using an exact Mixed Integer Linear Programming approach where the copy number of each plasmid bin is approximated using a network flow. For running gplas, we used the *E. coli* model of mlplasmids [5] to classify contigs as plasmidic or chromosomal prior to the binning step. We also used the classification probabilities computed by mlplasmids as the plasmid score for PlasBin-flow, keeping all other parameters with their default value. MOB-recon and HyAsP were run using their default parameters.

Ground truth plasmid bins were obtained for the 54 selected samples by mapping the short-read contigs to the hybrid contigs using BLAST+[9], discarding BLAST hits with identity below 95% or covering less than 80% of the short-read contig. For a given hybrid assembly contig (considered as a full plasmid), the corresponding ground truth plasmid bin is defined as the set of all short-read contigs belonging to hits to this hybrid contig.

The ground truth plasmid bins and the results from the four plasmid binning tools are available at https://zenodo.org/records/10785151.

In a practical context, the design of our experiments illustrate the use of PlasEval in a project where one would like to decide which plasmid binning tool to use in a specific project where both short read and long rad sequencing data is available for a set of isolates. A subset of samples for which hybrid sequencing data allow to recover a reasonable ground truth are used to benchmark a set of plasmid binning tools, and the PlasEval results allows to understand how these tools behave on this data and to select a tool to use on the whole dataset.

### 4.3 Evaluating accuracy against the ground truth

We first evaluated the results of all four binning methods using the notions of precision, recall and F1 score defined in Section 2. We then used PlasEval to compute the dissimilarity scores between the predicted plasmid bins of each plasmid binning tool and the ground truth. Figure 2 shows the correlation between the F1-score and the dissimilarity measure for all 54 samples.

**Fig. 2:**
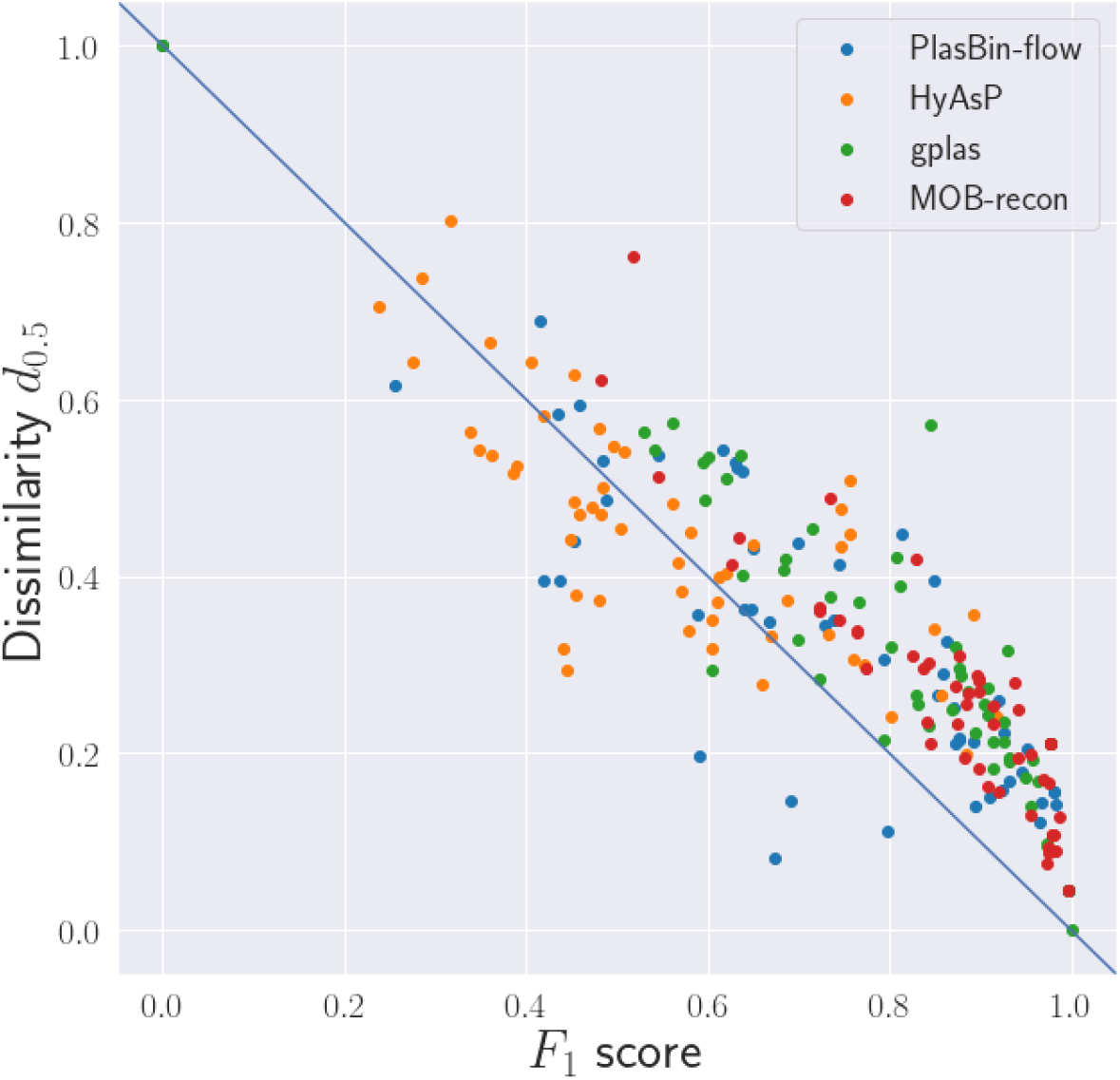
Correlation between F1 score and dissimilarity *D*

Despite the difference between the two accuracy measures, the dissimilarity score *d*_0.5_ is mostly inversely correlated to the F1 score: PlasBin-flow and HyAsP showed Pearson correlation coefficients of *−*0.85 and *−*0.76 respectively, while we observe a stronger correlation coefficient of *−*0.91 for the results of MOB-recon and gplas.

Overall this shows that the novel measure that we introduce is consistent with the F1 score. However, one of the interests of the dissimilarity measure is that it can be broken down into four components representing missing and extra contigs in predicted plasmid bins as well as the cost of splits and joins, thus providing a refined view of the errors in predicted plasmid bins. We illustrate this in Fig. 3 where we show the respective contribution of all four components of the dissimilarity measure *d*_0.5_ for five randomly chosen samples.

**Fig. 3:**
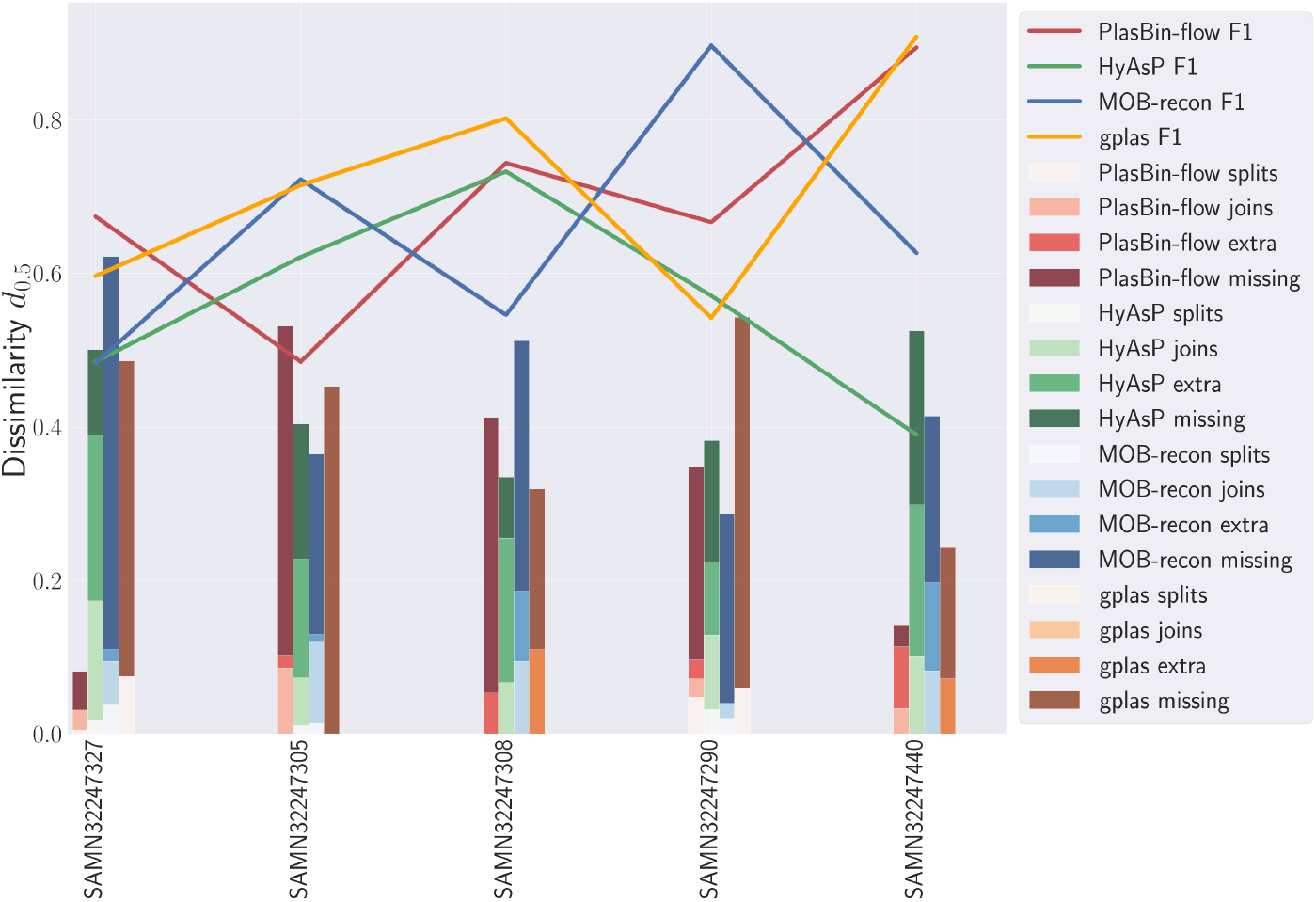
Dissimilarity comparison between all methods on five reference genome assemblies. The dissimilarity value is broken into its four components accounting for extra and missed contigs, splits and joins.

For instance, for sample SAMN32247327, we can see that MOB-recon and HyAsP have the same F1 scores but they have different dissimilarity *d*_0.5_ values and error types; the error in the plasmid bins predicted by gplas stem mostly from missing (plasmidic) contigs, while HyAsP errors are dominated by including in plasmid bins contigs not present in the ground truth (i.e. chromosomal contigs).

Table 1 below shows the mean and standard deviation of the F1-score and the PlasEval dissimilarity score for the four tools and over all samples. Supplementary Figure D.1 provides a detailed illustration of the contribution of the four components of the PlasEval score over all samples and all considered methods. Table 1 shows that overall, HyAsP is the least accurate of the three tools, and that the main errors in plasmid bins computed by HyAsP result from the inclusion in plasmid bins of chromosomal contigs. Interestingly, PlasBin-flow, a method based on a similar objective than HyAsP but using an exact optimization method instead of a greedy heuristic, is much more accurate than HyAsP; one can observe that its errors are mostly due to the omission of plasmidic contigs in the plasmid bins, while it is much more accurate than HyAsP for the other three components of the PlasEval score. This comparison illustrates the trade-off between an aggressive greedy heuristic (HyAsP) and an exact optimization method. Overall, the most accurate method on these samples is MOB-recon, followed by gplas. Similarly to PlasBin-flow, for both MOB-recon and gplas, the errors, as measured by the PlasEval score, stem from the omission of plasmidic contigs in the plasmid bins. Overall, more than 80% of the dissimilarity can be attributed to discrepancies in identifying plasmid contigs, suggesting that an avenue to improve plasmid binning is to improve true plasmidic contigs identification (contigs classification). Moreover, the patterns observed in terms of the contributions of splits and joins are different in the three most accurate methods PlasBin-flow, MOB-recon and gplas. For PlasBin-flow the contribution of splits is very low, while joins contribute more, suggesting it is a conservative method, that tends to not mix into a single plasmid bin contigs from different plasmids, at the expense of splitting true plasmids into several bins. The contribution of splits for gplas is somewhat high, suggesting a higher rate of true plasmids mixed into a single plasmid bin. Last, MOB-recon sits in-between with a low contribution of splits and a moderate contribution of joins.

**Table 1:** Mean and standard deviation of the PlasEval dissimilarity score and *F*_1_-score for the predictions of all four methods on 54 *E. coli* samples. The contribution of splits, joins, extra contigs and missing contigs to the dissimilarity is also shown.

| Method | Splits | Joins | Extra | Missing | $d_{0.5}$ | $F_1$ |
| --- | --- | --- | --- | --- | --- | --- |
| PlasBin-flow | **0.0018** ± **0.0076** | 0.0473 ± 0.0542 | **0.0464** ± **0.0484** | 0.2334 ± 0.1798 | 0.3299 ± 0.1879 | 0.7306 ± 0.2163 |
| HyAsP | 0.0051 ± 0.0113 | 0.0799 ± 0.0786 | 0.3000 ± 0.1821 | **0.0630** ± **0.0625** | 0.4479 ± 0.1333 | 0.5599 ± 0.1747 |
| gplas | 0.0148 ± 0.0310 | **0.0023** ± **0.0078** | 0.0459 ± 0.0553 | 0.2486 ± 0.1740 | 0.3116 ± 0.1727 | 0.8087 ± 0.1786 |
| MOB-recon | 0.0038 ± 0.0084 | 0.0198 ± 0.0354 | 0.0529 ± 0.0710 | 0.1776 ± 0.1218 | **0.2541** ± **0.1405** | **0.8684** ± **0.1263** |

The comparison of PlasBin-flow and gplas is interesting and illustrates well the use of PlasEval. Indeed, one can observe a similar overall score, similar contributions of missing and extra contigs, but a different pattern in the contributions of splits and joins. It follows that, if one is interested in choosing a plasmid binning tool that results in predicted plasmid bins that do not mix several true plasmids into a single bin, then PlasBin-flow is a more appropriate tool than gplas.

### 4.4 Comparing plasmid-binning tools

Alternatively to providing a refined measure of accuracy against a given ground truth, the PlasEval dissimilarity measure is useful to compare plasmid binning tools. Indeed, given two sets of predicted plasmid bins obtained with two different plasmid binning tools, an analysis similar to the one we conducted in the previous section can quantify the differences in the results of both tools.

Table 2 shows that with the exception of HyAsP, the dissimilarity between the other tools in terms of the predicted plasmid bins is generally found to be below 0.5. Most dissimilarity scores between the predictions of any two of PlasBin-flow, MOB-recon and gplas tend be between 0.1 and 0.5. When comparing the results of these tools against HyAsP, the dissimilarity scores generally range between 0.4 and 0.7. Figure 4 also shows that a small tail of high dissimilarity scores can be observed for almost all the pairs of tools.

**Table 2:** Mean and standard deviation for the different components of the dissimilarity score between results of different plasmid binning predictions.

| Method pairs | Splits | Joins | Extra | Missing | $d_{0.5}$ |
| --- | --- | --- | --- | --- | --- |
| PlasBin-flow vs. HyAsP | $0.0353 \pm 0.0433$ | $0.0196 \pm 0.0315$ | $0.0414 \pm 0.0576$ | $0.4311 \pm 0.1771$ | $0.5274 \pm 0.1526$ |
| PlasBin-flow vs. gplas | $0.0016 \pm 0.0075$ | $0.0912 \pm 0.0883$ | $0.0787 \pm 0.0707$ | $0.0745 \pm 0.0955$ | $0.2461 \pm 0.1300$ |
| PlasBin-flow vs. MOB-recon | $0.0111 \pm 0.0295$ | $0.0477 \pm 0.0662$ | $0.1127 \pm 0.0849$ | $0.1790 \pm 0.1711$ | $0.3505 \pm 0.1702$ |
| HyAsP vs. gplas | $0.0007 \pm 0.0038$ | $0.0670 \pm 0.0662$ | $0.4668 \pm 0.1500$ | $0.0458 \pm 0.0531$ | $0.5803 \pm 0.1346$ |
| HyAsP vs. MOB-recon | $0.0120 \pm 0.0218$ | $0.0716 \pm 0.0784$ | $0.4099 \pm 0.1555$ | $0.0421 \pm 0.0574$ | $0.5356 \pm 0.1355$ |
| gplas vs. MOB-recon | $0.0385 \pm 0.0516$ | $0.0030 \pm 0.0099$ | $0.0790 \pm 0.0829$ | $0.1568 \pm 0.1568$ | $0.2773 \pm 0.1773$ |

**Table 3:**
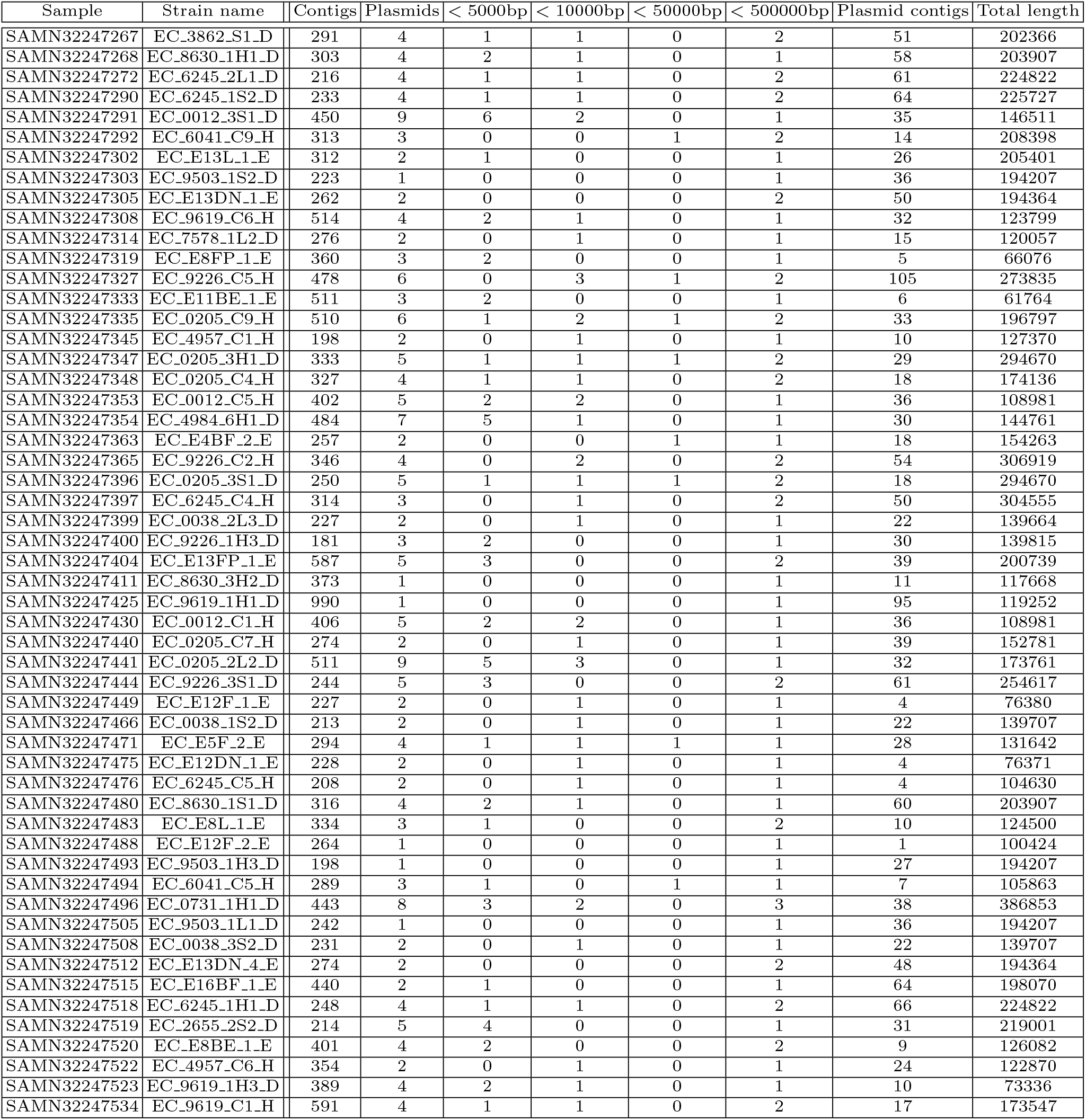
Statistics on input assemblies: number of contigs of the short-read assembly, number of plasmids from the hybrid assembly, number of plasmids by length (4 columns), number of short-read contigs in ground truth plasmid bins, total length of plasmids.

**Table 4:**
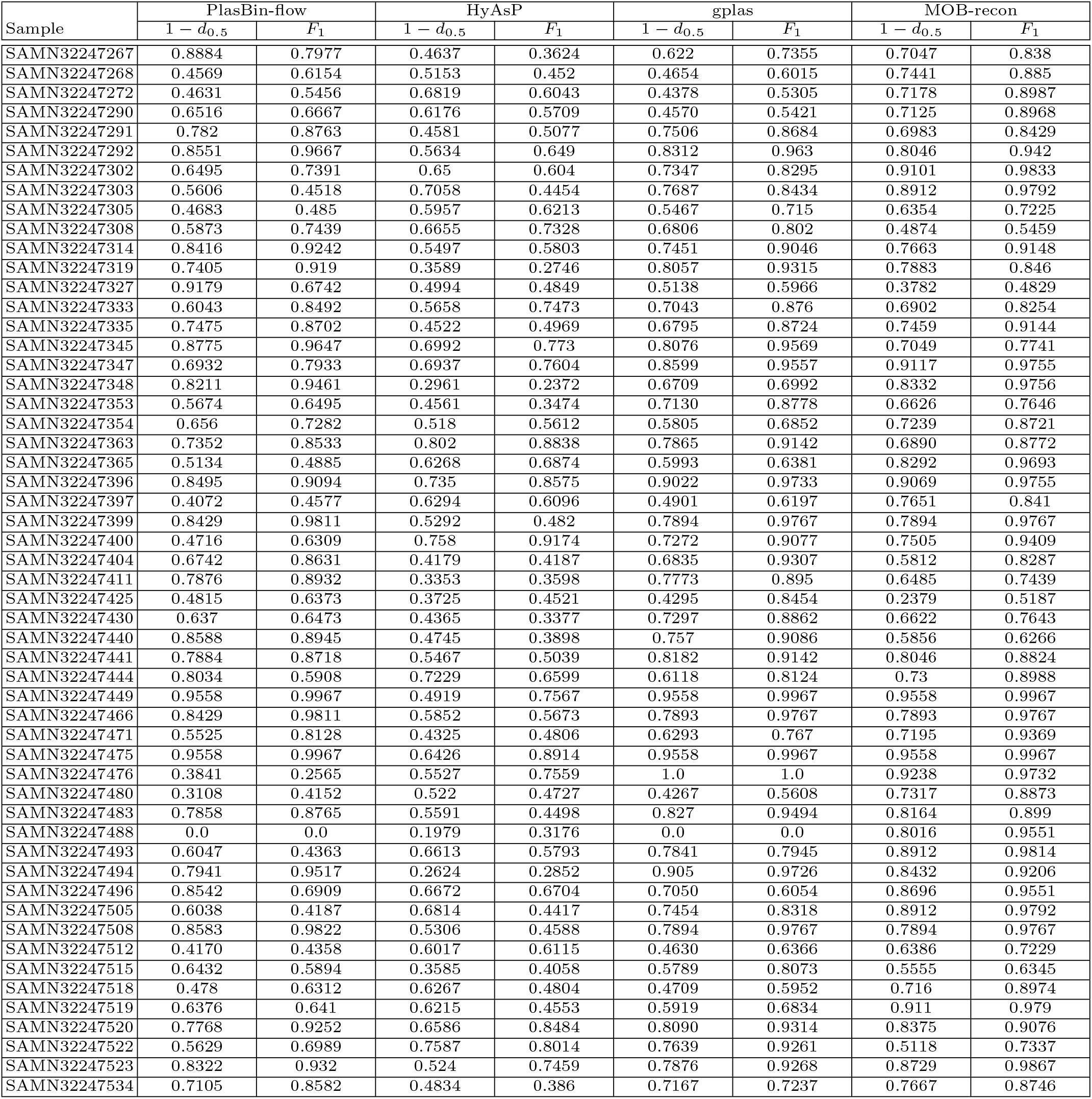
The *F*_1_-score and the dissimilarity score subtracted from 1, computed for the predictions of the 54 E. coli samples considered in this analysis.

**Fig. 4:**
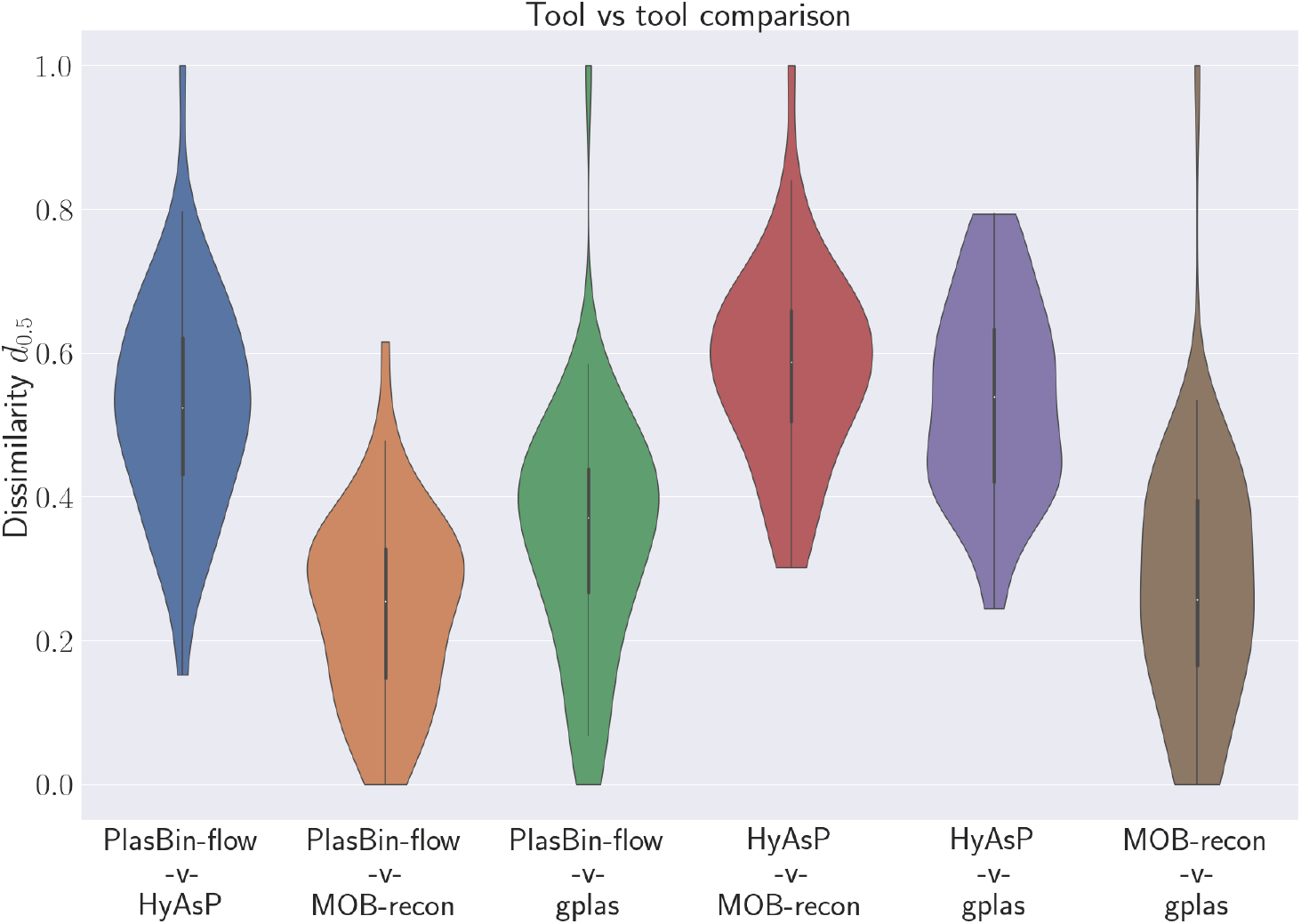
Distribution of the dissimilarity measure *d*_0.5_ between pairs of plasmid binning tools over all 54 genomes in the data set.

The two tools showing the largest agreement are PlasBin-flow and gplas. Joins (from PlasBin-flow to gplas) are the main source of dissimilarity between the two tools, indicating that plasmid bins from gplas tend to result from joining plasmid bins from PlasBin-flow. On the other hand, HyAsP predictions are highly dissimilar to those of other tools. This can be expected since PlasBin-flow, gplas and MOB-recon seem to have a relatively conservative approach to selecting plasmid contigs as compared to HyAsP. Compared against MOB-recon and HyAsP, PlasBin-flow excludes in its plasmid bins a large number of contigs that the other methods contain in their predictions. More interestingly, we observe that differences in plasmid bins are mostly due to plasmid bins of PlasBin-flow that are joined in MOB-recon. Last, when comparing MOB-recon and gplas, we observe that both methods differ in terms of contigs that appear only in bins of one method, followed by the fact that a significant number of plasmid bins from MOB-recon occur in a single gplas bin, while to the contrary, gplas bins do not exhibit such a feature. Supplementary Figures E.2 and E.3 provide a refined illustration of the differences discussed above.

## 5 Conclusion

In this work, we introduce a framework that enables to compare two sets of predicted plasmid bins. We provide a measure to quantify the disagreement between the two sets through a dissimilarity score, whose individual components (missing and extra contigs, splits, joins) provide both a quantitative and qualitative assessment of the dissimilarity between the two considered sets of plasmid bins. Moreover, the introduced dissimilarity measure accounts for important features such as contig length and copy number. This is especially important as common repeats, such as Insertion Sequences (IS) are known to be widely present in plasmids.

We applied this method for evaluating the predicted plasmids from four methods (PlasBin-flow, HyAsP, gplas and MOB-recon) on a data set of 54 *E. coli* samples for which the ground truth plasmid bins were obtained from hybrid assemblies. Our analysis shows that PlasEval provides a useful way to compare predicted plasmid bins either against a ground truth or between pairs of plasmid binning tools, that can inform either developers of novel plasmid binning methods or potential users having to decide on using a specific plasmid binning tool on a specific dataset. The results we obtained when analyzing the predictions of gplas, HyAsP, MOB-recon and PlasBin-flow show that the methods do behave significantly differently. This suggests that the problem of plasmid binning from short-reads data is still in need of further developments and we believe that PlasEval will be a useful tool toward this effort.

## Acknowledgments

This research was funded by Genome Canada and NSERC.

## A Proofs and additional lemmas

*Proof of Lemma 1.* 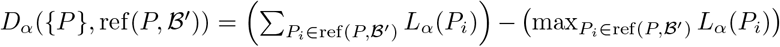 It can be easily shown that ref(*P, ℬ*′) is the optimal way to partition *P*. There no unique contigs between *P* and ref(*P, ℬ*′). Say *P* = *P*_1_ *∪ P*_2_ *∪* … *∪ P*_*k*_ such that *L*_*α*_(*P*_1_) *≤ L*_*α*_(*P*_2_) *≤* … *≤ L*_*α*_(*P*_*k*_).

While partitioning any set into two sets, the size of the smaller set counts towards the cost of the split. Thus, we repeatedly partition *P* to obtain ref(*P, ℬ*′) by choosing the smallest set in ref(*P, ℬ*′) to split first.

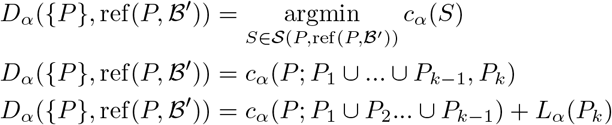

Repeatedly removing the smallest set of ref(*P, ℬ*′) available, we get

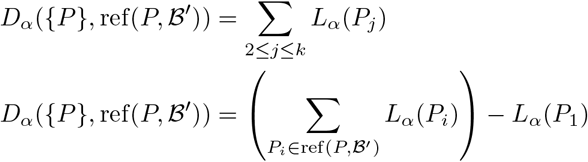

Since, *P*_1_ was the set in ref(*P, ℬ*′) with the largest size, this proves the lemma.

We now introduce a lemma that implies that, when comparing two sets of bins *𝒜, ℬ*, adding a contig *c* of length *ℓ* to a bin *P* ∈ 𝒜 and *Q* ∈ *ℬ*, will not result in either a decrease of the most parsimonious cost of the splits transforming *𝒜* into ref(*𝒜*′, *ℬ*′) (and, by symmetry between splits and joins, of the joins transforming *ℬ*′ ref(*𝒜*′, *ℬ*′)) or an increase of the cost of these splits (resp. joins) of more than *ℓ* ^*α*^. We will use this Lemma in the bounding function for the branch-and-bound algorithm we define to handle the case of repeated contigs.

### Lemma 2.

*Let P be a plasmid bin, P*_0_, *P*_1_, …, *P*_*k*_ *be such that P*_0_ = *∅ and* 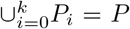. *Let c be a contig not in P, of length ℓ. Let* 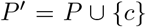 *and* 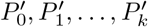 *such that (1) there exists j with* 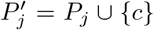 *and (2) for any* 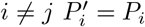 *Then*

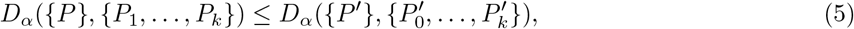

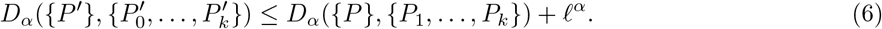

*Proof*.

First, as *P*_0_ = *∅*, it does not contribute anything to *D*_*α*_({*P }, {P*_0_, …, *P*_*k*_}), as *L*(*P*_0_) = 0, which implies that *D*_*α*_({*P }, {P*_1_, …, *P*_*k*_}) = *D*_*α*_({*P }, {P*_0_, …, *P*_*k*_}).

For similar reasons, if *j≠*0, then 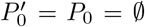 so 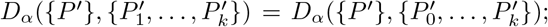 the only case where this does not hold is when 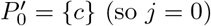.

Next, without loss of generality, assume that for any *i, L*(*P*_*i*_) *≤ L*(*P*_*k*_), i.e. *P*_*k*_ is the subset of *P* of maximal cumulative length. It follows from Lemma 1 that

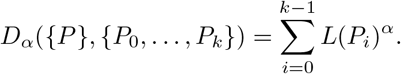

We first assume that *L*(*P* ′) *> L*(*P*_*k*_). Lemma 1 implies that

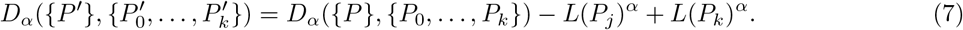

As *L*(*P*_*k*_) *< L*(*P*_*j*_) + *𝓁* and *α ≥* 0, we have that *L*(*P*_*k*_)^*α*^ *≤* (*L*(*P*_*j*_) + *𝓁*)^*α*^. Moreover, as *α ≤* 1, (*L*(*P*_*j*_) + *𝓁*)^*α*^ *≤ L*(*P*_*j*_)^*α*^ + *𝓁*^*α*^. This implies that *L*(*P*_*k*_)^*α*^ *< L*(*P*_*j*_)^*α*^ + *𝓁*^*α*^, which, combined with (7), implies (6). Moreover, as *L*(*P*_*k*_) *≥ L*(*P*_*j*_), *L*(*P*_*k*_)^*α*^ *≥ L*(*P*_*j*_)^*α*^, which, combined with (7), implies (5).

We now assume that 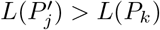 Lemma 1 implies that

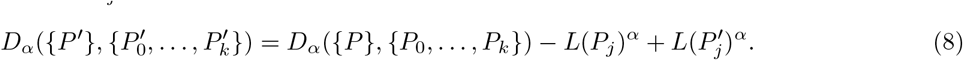

As 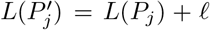 we have 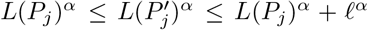 which, combined with (8), implies (6). Moreover, as 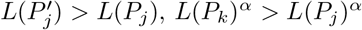, which, combined with (8), implies (5).

## B Branch-and-bound algorithm

The branch-and-bound algorithm searches the space *ℳ* (*𝒜, ℬ*) of all matchings between the copies of contigs of *𝒜, ℬ* to find a matching with the optimal dissimilarity score. It explores a search tree that is constructed by introducing a contig family containing repeats per level.

The root is at level 0, where we have *𝒜*_0_, *ℬ*_0_ consisting of only non-repeat contig families. As a result, the matching at this level is unambiguous. Each subsequent level introduces a contig family containing repeated contigs. For the family of a particular contig *c*, if *𝒜, ℬ* have *k*_1_, *k*_2_ copies respectively, the total number of possible matchings for this family is 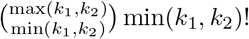 Each edge from a search tree node at level *i −* 1 to one at level *i* represents a choice of a specific matching between the contig copies for the family at level *i*. Once a matching is chosen, we resolve the matching for the contig family *c*, while the remaining |*k*_1_ *− k*_2_| copies of *c* are removed as unique copies.

Note that before resolving the matching for a contig family at level *i*, the matchings for contig families at levels 1 to *i −* 1 have already been resolved. As a result, each node of the search tree depicts a unique matching of copies of contigs from families at levels 1 to *i −* 1 as well as the contigs from non-repeat families from level 0. At each tree search node, after resolving the matching for all contig families at the node, we compute the ref(*P, ℬ*′)_*i*_ for each *P* ∈ 𝒜′. This is the refinement of *P* according to bins in *ℬ*′ while only accounting for contig families encountered till level *i*. This helps in computing the splits to be made from 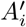. Similarly, we can compute the joins to be made from ref(*𝒜*′, *ℬ*′)_*i*_ to *ℬ*_*i*_.

### Algorithm 1

Branch and bound algorithm for transforming *𝒜* to *ℬ*

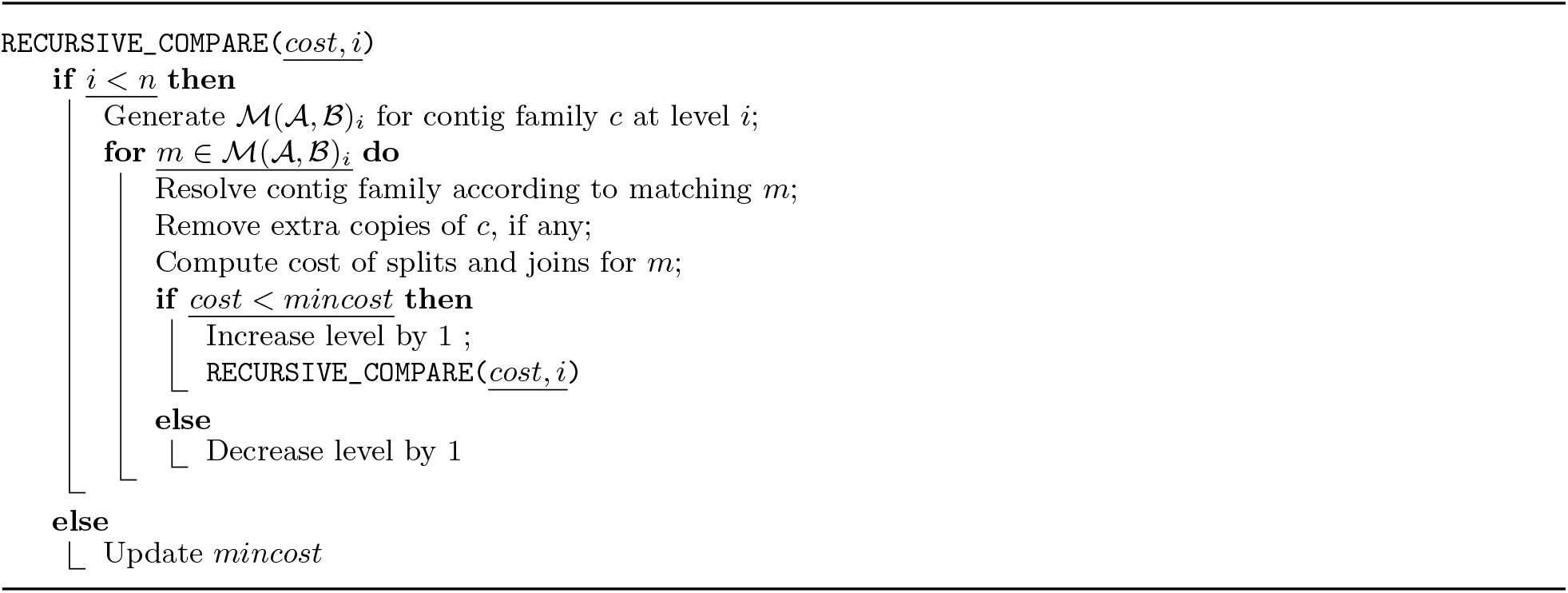

## C Data

## D Tool evaluations using PlasEval

**Fig. D.1:**
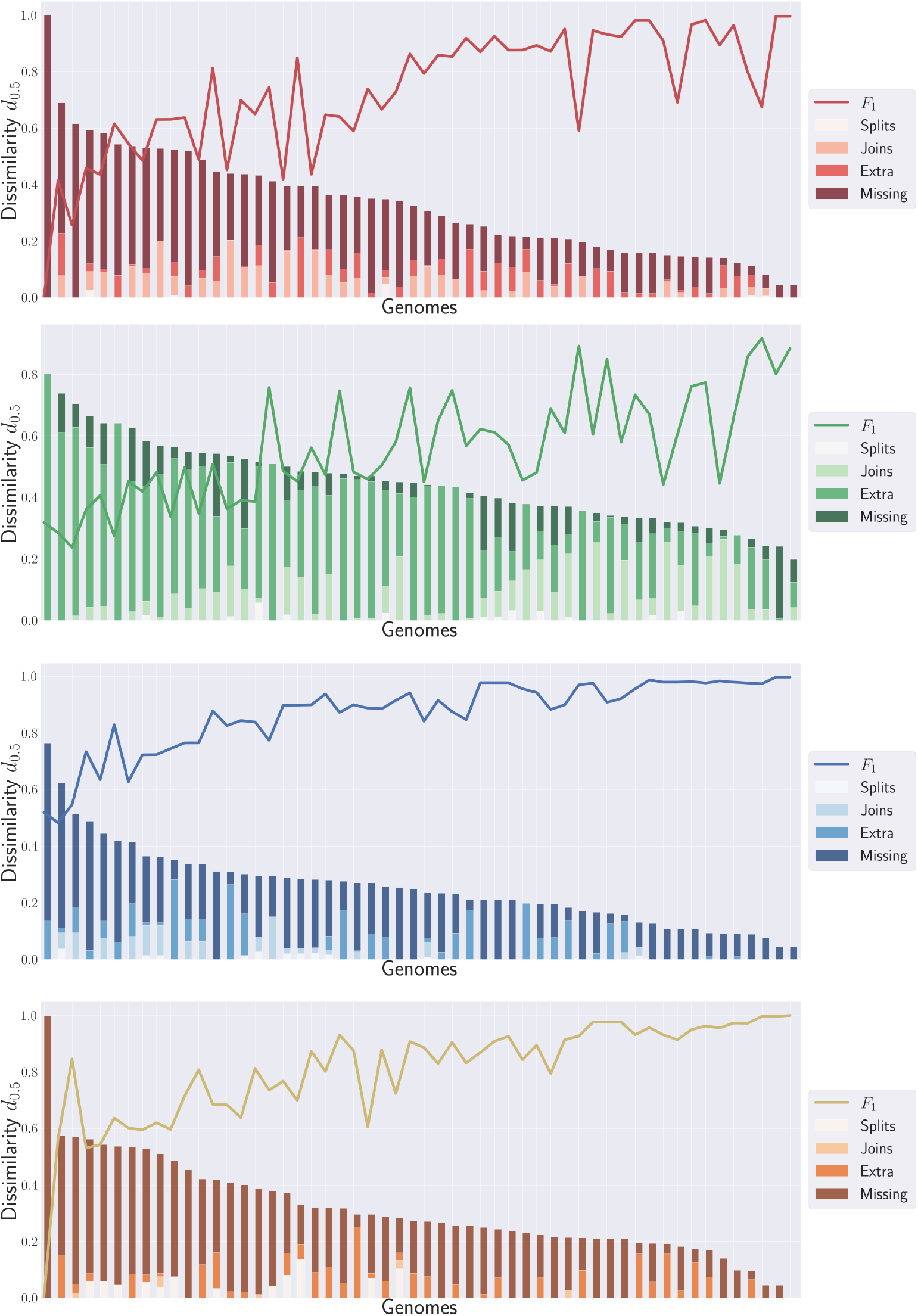
PlasEval score and F1 score for all samples and all tools. Genomes on the x-axis are sorted in decreasing order of dissimilarity with respect to the reference. From top to bottom: PlasBin-flow, HyAsP, MOB-recon, gplas.

## E Tool versus tool comparisons

Figures E.2 and E.3 provide further details on the differences between the binning predictions of PlasBin-flow, MOB-recon, gplas and HyAsP.

**Fig. E.2:**
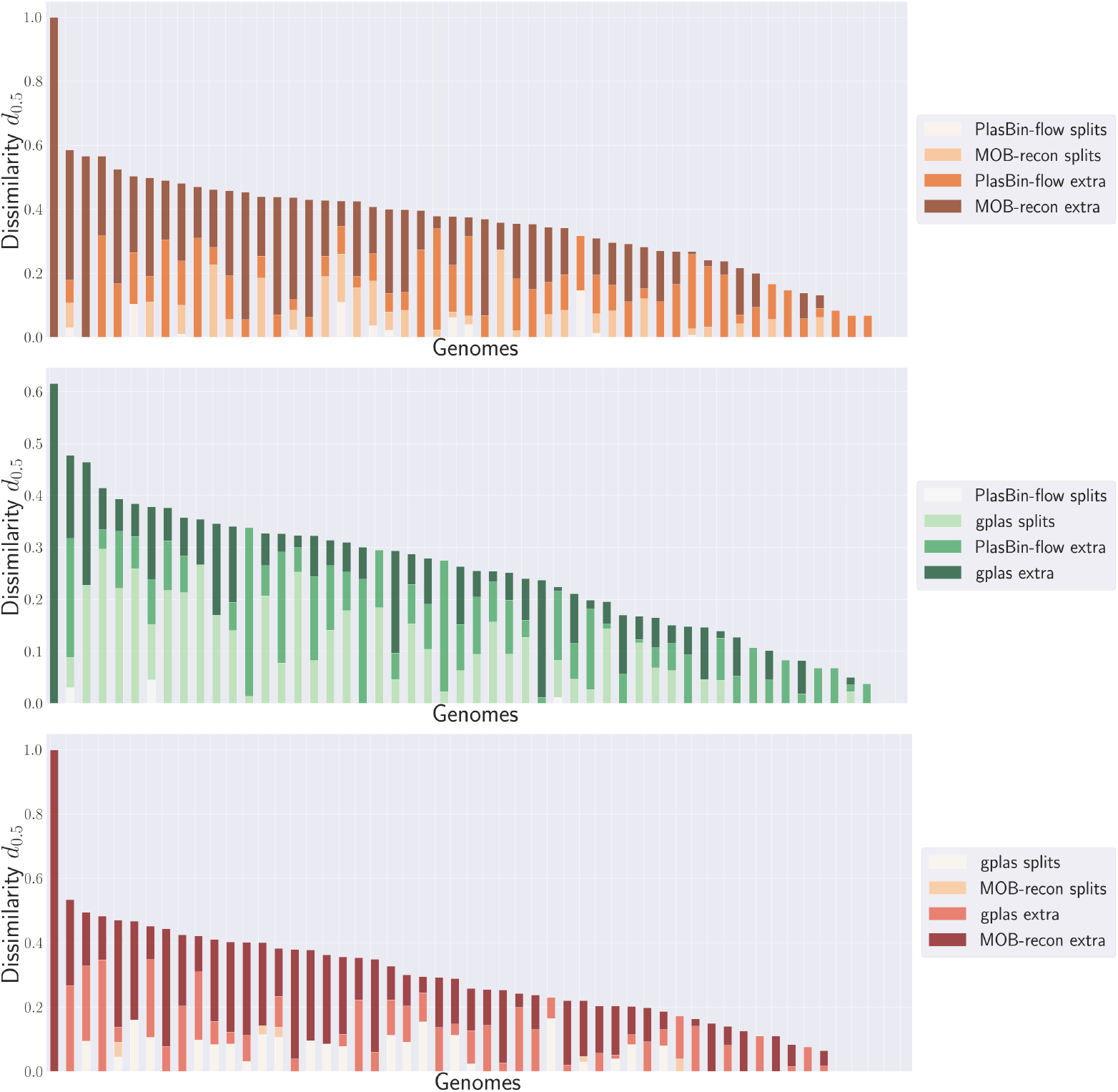
Pairwise comparisons between PlasBin-flow, MOB-recon and gplas.

**Fig. E.3:**
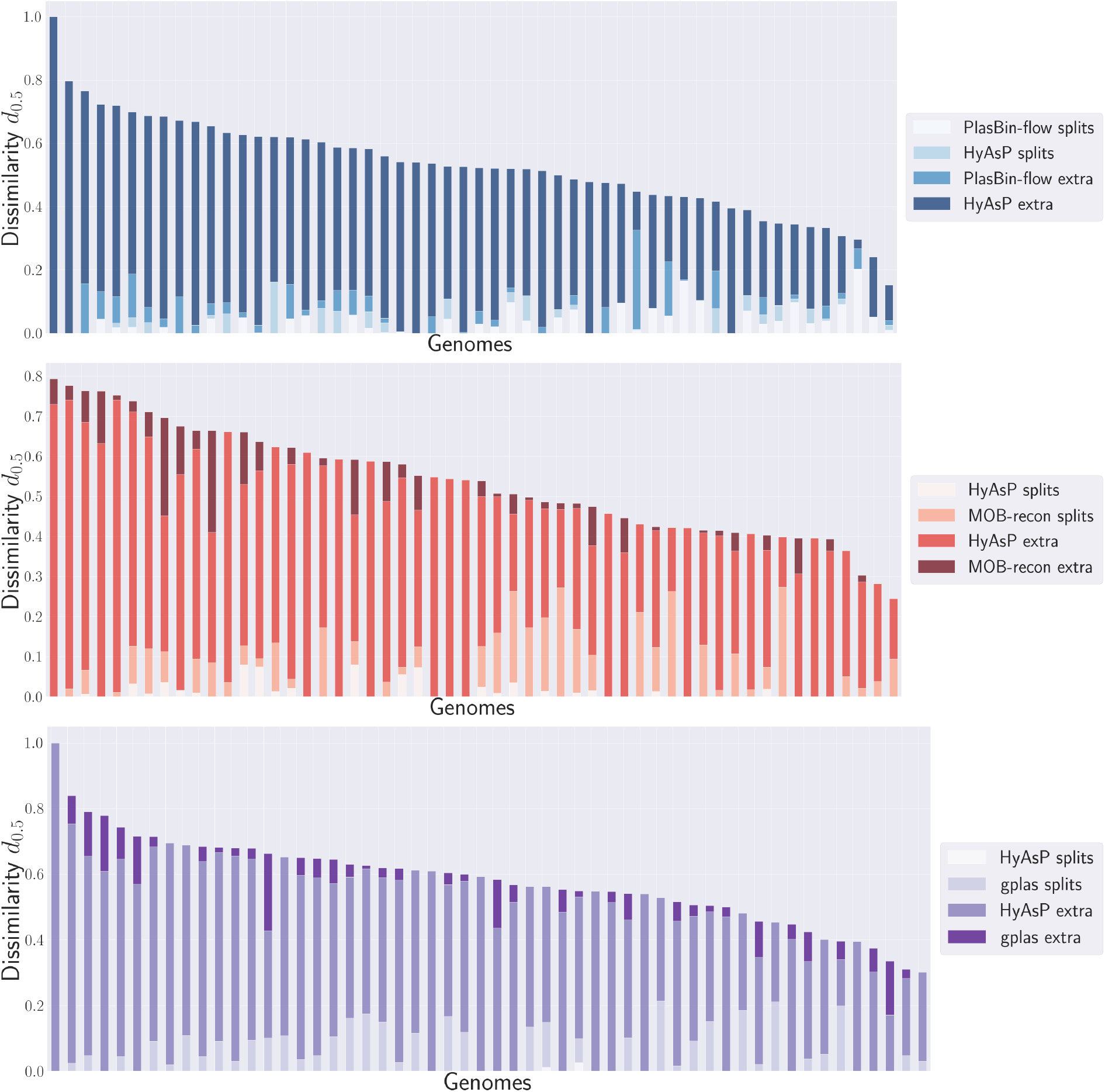
Pairwise comparisons between HyAsP and PlasBin-flow, MOB-recon and gplas.

## F Running time and function calls

| Comparison | Mean time (s) | Max time (s) | Mean # calls | Max # calls |
| --- | --- | --- | --- | --- |
| PlasBin-flow - Ground truth | 0.18 | 6.81 | 545 | 19673 |
| HyAsP - Ground truth | 1.14 | 52.63 | 980 | 40105 |
| Gplas - Ground truth | 0.002 | 0.02 | 9 | 24 |
| MOB-recon - Ground truth | 0.003 | 0.02 | 11 | 28 |
| PlasBin-flow - HyAsP | 0.43 | 15.28 | 658 | 24459 |
| PlasBin-flow - Gplas | 0.005 | 0.1 | 22 | 400 |
| PlasBin-flow - MOB-recon | 0.27 | 1.28 | 95 | 4368 |
| HyAsP - Gplas | 0.003 | 0.009 | 10 | 23 |
| HyAsP - MOB-recon | 0.004 | 0.024 | 11 | 28 |
| Gplas - MOB-recon | 0.001 | 0.01 | 8 | 16 |

## Notes

https://github.com/acme92/PlasEval

### Competing Interest Statement

The authors have declared no competing interest.

## References

1. Andreopoulos, W.B., Geller, A.M., Lucke, M., Balewski, J., Clum, A., Ivanova, N.N., Levy, A.: Deeplasmid: deep learning accurately separates plasmids from bacterial chromosomes. Nucleic Acids Research 50(3), e17–e17 (12 2021). 10.1093/nar/gkab1115, https://doi.org/10.1093/nar/gkab1115

2. Andrews, S.: FastQC: a quality control tool for high throughput sequence data (2010), http://www.bioinformatics.babraham.ac.uk/projects/fastqc

3. Antipov, D., Hartwick, N., Shen, M., Raiko, M., Lapidus, A., Pevzner, P.A.: plasmidSPAdes: assembling plasmids from whole genome sequencing data. Bioinformatics 32(22), 3380–3387 (2016). 10.1093/bioinformatics/btw493

4. Arredondo-Alonso, S., Bootsma, M., Hein, Y., Rogers, M.R.C., Corander, J., Willems, R.J.L., Schürch, A.C.: gplas: a comprehensive tool for plasmid analysis using short-read graphs. Bioinformatics 36(12), 3874–3876 (2020). 10.1093/bioinformatics/btaa233

5. Arredondo-Alonso, S., Rogers, M.R.C., Braat, J.C., Verschuuren, T.D., Top, J., Corander, J., Willems, R.J.L., Schürch, A.C.: mlplasmids: a user-friendly tool to predict plasmid- and chromosome-derived sequences for single species. Microbial Genomics 4(11), e000224 (2018). 10.1099/mgen.0.000224

6. Arredondo-Alonso, S., Willems, R.J., van Schaik, W., Schürch, A.C.: On the (im)possibility of reconstructing plasmids from whole-genome short-read sequencing data. Microbial Genomics 3(10) (2017). 10.1099/mgen.0.000128

7. van der Graaf-van Bloois, L., Wagenaar, J.A., Zomer, A.L.: Rfplasmid: predicting plasmid sequences from short-read assembly data using machine learning. Microbial Genomics 7(11), 000683 (2021). 10.1099/mgen.0.000683, https://www.microbiologyresearch.org/content/journal/mgen/10.1099/mgen.0.000683

8. Bouras, G., Sheppard, A.E., Mallawaarachchi, V., Vreugde, S.: Plassembler: an automated bacterial plasmid assembly tool. Bioinformatics 39(7) (2023). 10.1093/bioinformatics/btad409

9. Camacho, C., Couluris, G., Avagyan, V., Ma, N., Papadopoulos, J., Bealer, K., Madden, T.L.: BLAST+: architecture and applications. BMC Bioinformatics 10, 421 (2009). 10.1186/1471-2105-10-421

10. Chen, S., Zhou, Y., Chen, Y., Gu, J.: fastp: an ultra-fast all-in-one fastq preprocessor. Bioinfotmatics 34(17), i884–i890 (2018). 10.1093/bioinformatics/bty560

11. Chen, Z., Erickson, D.L., Meng, J.: Benchmarking hybrid assembly approaches for genomic analyses of bacterial pathogens using Illumina and Oxford Nanopore sequencing. BMC Genomics 21(1) (2020). 10.1186/s12864-020-07041-8

12. Dasgupta, B., Jiang, T., Kannan, S., Li, M., Sweedyk, E.: On the complexity and approximation of syntenic distance. Discrete Applied Mathematics 88(1–3), 59–82 (1998). 10.1016/s0166-218x(98)00066-3

13. De Oliveira, D.M.P., Forde, B.M., Kidd, T.J., Harris, P.N.A., Schembri, M.A., Beatson, S.A., Paterson, D.L., Walker, M.J.: Antimicrobial resistance in ESKAPE pathogens. Clinical Microbiology Reviews 33(3) (2020). 10.1128/cmr.00181-19

14. Dusadeepong, R., Delvallez, G., Cheng, S., Meng, S., Sreng, N., Letchford, J., Choun, K., Teav, S., Hardy, L., Jacobs, J., Hoang, T., Seemann, T., Howden, B.P., Glaser, P., Stinear, T.P., Vandelannoote, K.: Phylogenomic investigation of an outbreak of fluoroquinolone-resistant salmonella enterica subsp. enterica serovar paratyphi a in phnom penh, cambodia. Microbial Genomics 9(3) (2023). 10.1099/mgen.0.000972

15. Ferretti, V., Nadeau, J.H., Sankoff, D.: Original synteny. In: Combinatorial Pattern Matching, 7th Annual Symposium, CPM 96. Lecture Notes in Computer Science, vol. 1075, pp. 159–167. Springer (1996). 10.1007/3-540-61258-0_13

16. Gurevich, A., Saveliev, V., Vyahhi Nikolayand Tesler, G.: Quast: quality assessment tool for genome assemblies. Bioinfotmatics 29(8), 1072–1075 (2013). 10.1093/bioinformatics/btt086

17. Johnson, J., Soehnlen, M., Blankenship, H.M.: Long read genome assemblers struggle with small plasmids. Microbial Genomics 9(5) (2023). 10.1099/mgen.0.001024

18. Khezri, A., Avershina, E., Ahmad, R.: Hybrid assembly provides improved resolution of plasmids, antimicrobial resistance genes, and virulence factors in escherichia coli and klebsiella pneumoniae clinical isolates. Microorganisms 9(12), 2560 (2021). 10.3390/microorganisms9122560

19. Mane, A., Faizrahnemoon, M., Vinař, T., Brejová, B., Chauve, C.: PlasBin-flow: a flow-based MILP algorithm for plasmid contigs binning. Bioinformatics 39(Supplement 1), i288–i296 (2023). 10.1093/bioinformatics/btad250

20. Mane, A.C., Faizrahnemoon, M., Chauve, C.: A mixed integer linear programming algorithm for plasmid binning. In: Jin, L., Durand, D. (eds.) Comparative Genomics - 19th International Conference, RECOMB-CG 2022, La Jolla, CA, USA, May 20-21, 2022, Proceedings. Lecture Notes in Computer Science, vol. 13234, pp. 279–292. Springer (2022). 10.1007/978-3-031-06220-9_16

21. Mikheenko, A., Prjibelski, A., Saveliev, V., Antipov, D., Gurevich, A.: Versatile genome assembly evaluation with QUAST-LG. Bioinformatics 34(13), i142–i150 (2018). 10.1093/bioinformatics/bty266

22. Müller, R., Chauve, C.: HyAsP, a greedy tool for plasmids identification. Bioinformatics 35(21), 4436–4439 (2019). 10.1093/bioinformatics/btz413

23. Partridge, S.R., Kwong, S.M., Firth, N., Jensen, S.O.: Mobile genetic elements associated with antimicrobial resistance. Clinical Microbiology Reviews 31(4) (2018). 10.1128/cmr.00088-17

24. Pradier, L., Tissot, T., Fiston-Lavier, A., Bedhomme, S.: Plasforest: a homology-based random forest classifier for plasmid detection in genomic datasets. BMC Bioinform. 22(1), 349 (2021). 10.1186/S12859-021-04270-W, https://doi.org/10.1186/s12859-021-04270-w

25. Robertson, J., Nash, J.: MOB-suite: software tools for clustering, reconstruction and typing of plasmids from draft assemblies. Microbial Genomics 4 (07 2018). 10.1099/mgen.0.000206

26. Rozov, R., Brown Kav, A., Bogumil, D., Shterzer, N., Halperin, E., Mizrahi, I., Shamir, R.: Recycler: an algorithm for detecting plasmids from de novo assembly graphs. Bioinformatics 33(4), 475–482 (2016). 10.1093/bioinformatics/btw651

27. Sanderson, H., Gray, K.L., Manuele, A., Maguire, F., Khan, A., Liu, C., Navanekere Rudrappa, C., Nash, J.H.E., Robertson, J., Bessonov, K., Oloni, M., Alcock, B.P., Raphenya, A.R., McAllister, T.A., Peacock, S.J., Raven, K.E., Gouliouris, T., McArthur, A.G., Brinkman, F.S.L., Fink, R.C., Zaheer, R., Beiko, R.G.: Exploring the mobilome and resistome of Enterococcus faecium in a One Health context across two continents. Microbial Genomics 8(9) (2022). 10.1099/mgen.0.000880

28. Sanderson, H., Nnajide, C., McCarthy, M., Singh, R., Rubin, J., Dillon, J.A., White, A.: Hybrid Genome Assemblies of 245 Avian and Broiler Barn Environment-Associated Escherichia coli Strains Isolated from Saskatchewan Broiler Farms. Microbiology resource announcements 12, e0011023 (04 2023). 10.1128/mra.00110-23

29. Shintani, M., Sanchez, Z.K., Kimbara, K.: Genomics of microbial plasmids: classification and identification based on replication and transfer systems and host taxonomy. Frontiers in Microbiology 6 (2015). 10.3389/fmicb.2015.00242

30. Sielemann, J., Sielemann, K., Brejova, B., Vinař, T., Chauve, C.: plASgraph2: using graph neural networks to detect plasmid contigs from an assembly graph. Frontiers in Microbiology 14 (2023). 10.3389/fmicb.2023.1267695

31. Wick, R.R., Judd, L.M., Gorrie, C.L., Holt, K.E.: Unicycler: Resolving bacterial genome assemblies from short and long sequencing reads. PLOS Computational Biology 13(6), 1–22 (06 2017). 10.1371/journal.pcbi.1005595

